# The Impact of Delayed Evacuation on the Quality of Human Fetal Tissue

**DOI:** 10.1101/2025.07.15.663318

**Authors:** Yasmeen Otaibi, Kevin Lee, Lucinda Cort, Diana O’Day, Jennifer C. Dempsey, Ian Phelps, Elizabeth Micks, Lyndsey S. Benson, Mei Deng, Dan Doherty, Sarah Prager, BDRL, Ian A. Glass, Kimberly A. Aldinger

## Abstract

**Background:** Digoxin and other agents are frequently administered to arrest fetal circulation before the evacuation of human fetal tissue (HFT) in pregnancy terminations to address concerns about fetal viability. However, the impact of delayed evacuation on HFT quality remains unknown. Analyzing HFT is critical for diagnosing pregnancies affected by fetal abnormalities and for driving progress in biomedical research.

**Objective:** This study aims to assess the effects of delayed evacuation following fetal circulation cessation on the quality of HFT for both diagnostic and research purposes.

**Study Design:** HFT samples were collected from second-trimester dilation and evacuation (D&E) procedures, with and without agent injection approximately 24 hours prior, as per standard care protocols. We assessed multiple parameters relevant to diagnostics and research, including: 1) cell morphology, 2) cell proliferation, 3) apoptosis, 4) cell viability in culture, and 5) nucleic acid quality. To simulate in utero conditions and determine the timeline for tissue degradation, we incubated brain tissue obtained from D&E without induced demise at 37°C for up to 18 hours.

**Results:** We analyzed 18 HFT samples from D&E procedures performed 19-25 hours after digoxin or potassium chloride (KCl) injection and compared them to 26 HFT samples from immediate evacuations without prior injection. Induced fetal demise resulted in 1) disrupted cell morphology, 2) decreased cell proliferation, 3) increased apoptosis, 4) reduced cell viability in culture, and 5) lower RNA quality. Despite these findings, all samples yielded DNA of sufficient quality for polymerase chain reaction (PCR).

**Conclusion:** D&E procedures performed after fetal demise induced by digoxin or KCl lead to decreased HFT quality, limiting its diagnostic and research potential beyond gross tissue evaluation and DNA extraction. Limiting the time between fetal circulation arrest and evacuation may improve HFT quality for clinical and research applications.

## INTRODUCTION

Evaluation of fetal tissue can be critical for determining a specific biomedical diagnosis in pregnancies affected by fetal abnormalities. A specific diagnosis is essential for accurate genetic counseling of recurrence risk and outcomes for future pregnancies. The historical standard for fetal diagnosis is an autopsy by a fetal pathologist, which can include radiological examinations, gross and microscopic tissue evaluation (including immunohistochemical staining), cellular assays, and select biochemical testing.^1,2^ Increasingly, fetal DNA is used for microarray testing to detect pathogenic chromosome copy number variants, as well as single gene, gene panel, and exome or whole genome sequencing (WGS) approaches to identify causative DNA variants.^2–5^

In parallel with clinical applications, HFT obtained after pregnancy termination has played a vital role in research advances that have greatly improved health care over decades. Human fetal-derived cells used for viral culture were essential for generating vaccines to poliovirus, varicella, rubella, measles and other pathogens.^6–10^ Vaccines created using the WI-38 fetal lung cell line alone are estimated to have averted or ameliorated an estimated 4.5 billion cases of poliomyelitis, measles, mumps, rubella, varicella, herpes zoster, adenovirus, rabies and Hepatitis A infection worldwide from 1960 to 2015.^11^ The human embryonic kidney293 (*HEK293*) fetal cell line has been widely used in a number of studies, including recombinant protein production, gene therapy, and drug development, as well as more recently in the development of RNA-based vaccines for COVID-19.^13,14,15^ HFT provision for the Human Lung Cell Atlas has also facilitated insights into the precise pathogenesis of COVID-19 infection.^12^ Furthermore, innumerable benefits of HFT provision to many other biomedical research efforts exist including developing immunotherapies for cancer, stem cell biology, tissue and organ cell regeneration, human disease modeling in animal models and characterizing birth defects, among others.^16^

In the United States, most second trimester pregnancy terminations are completed by dilation and evacuation (D&E).^17,18^ D&E procedures are often preceded by the use of digoxin, potassium chloride (KCl), and other agents to arrest fetal circulation.^19^ This approach is used for both termination of pregnancies (TOP) with structurally normal fetuses and those with fetal anatomic anomalies, genetic disorders, infections, or teratogenic exposures.^19^ While no evidence-based medical benefits of this practice have been demonstrated,^20^ some clinicians perceive increased procedural ease and safety.^21^ In addition, institution, provider, and/or patient preference may play a role. In the U.S., clinician concern for legal ramifications of the Partial Birth Abortion Ban Act of 2003 is also influential.^19,22^

Methods of inducing fetal demise include injecting the agent into the fetus, umbilical cord, or amniotic fluid, typically followed by D&E procedure or labor induction.^19^ While concerns have been raised that the method of termination may adversely impact the utility of fetal tissues,^23^ no studies have systematically evaluated the effects of agents that arrest fetal circulation or the timing of D&E after injection on HFT quality. To address this critical knowledge gap, the goal of this study was to evaluate HFT quality using a variety of parameters that evaluate utility of these tissues for diagnostic and research purposes.

## MATERIALS AND METHODS

### Collection of Human Fetal Tissue

Specimens were obtained from several outpatient reproductive health clinics in the Seattle area between March 18^th^, 2014 and December 7^th^, 2018. All specimens were collected from pregnant participants who chose to donate HFT during a D&E procedure. Second-trimester pregnancy terminations comprised most donations, with four out of 44 specimens being early third-trimester. Written informed consent was obtained from all participants under a protocol approved by the University of Washington Institutional Review Board (STUDY00000380).

Participants were consented broadly through the Birth Defect Research Laboratory (BDRL) for potential use of donated tissue in biomedical research only after consent for the procedure was obtained. Clinics followed varying protocols concerning the gestational duration at which digoxin or KCl was administered, with most using these agents after 18-20 weeks post- fertilization.

HFT from 44 specimens were included in this study (Table S1). HFT was collected from D&E procedures performed approximately 24 hours (19-25 hours) after the induction of fetal demise using digoxin (intra-fetal or intra-amniotic) or KCl (intra-cardiac or intra-funic), consistent with standard care practices in Washington State. Injections coincided with the placement of cervical dilators per clinic protocols. Providers confirmed fetal demise via ultrasound prior to the D&E. Specimens were also collected from D&E procedures completed without prior induction of fetal demise. Tissues that were not processed immediately were stored at 4°C during transport. Upon arrival to the laboratory, all tissues were briefly washed in Hanks’ balanced salt solution (HBSS) prior to processing for downstream applications. Participant data was de-identified and assigned a study ID number to be analyzed anonymously. The gestational duration of the fetus was estimated from fetal foot length and reported as the number of days post-fertilization.^24^ We compared tissues from specimens evacuated 24 hours after digoxin or KCl injection (delayed evacuation) to tissues from specimens evacuated without prior injection (immediate evacuation). Sample size calculations were not performed for this descriptive study, and not all specimens were sufficiently intact to provide HFT from all target organs (brain, kidney, liver, lung, muscle).

The number of specimens for both immediate and delayed evacuation was insufficient for consistent testing across all evaluations: histological evaluations (3 delayed and 3 immediate evacuations), cell viability tests (3 delayed and 7 immediate evacuations), cell culture tests (11 delayed and 9 immediate evacuations), and nucleic acid tests (8 delayed and 2 immediate evacuation specimens).

To assess the time course of tissue degradation following the cessation of fetal circulation, we simulated in utero temperatures by incubating tissues from two immediate evacuation specimens at 37°C in Gibco Hibernate-E for 6 and 18 hours. Incubated control specimens were analyzed as described below and compared to delayed evacuation tissues and non-incubated immediate evacuation tissues. To investigate the potential effects of digoxin on tissue quality, we incubated tissues from four immediate evacuation specimens in Gibco Hibernate-E with and without digoxin (1 mg/500 mL media) at both 37°C and 4°C for 18 hours.

### Histological Evaluation & Cell Culture

Brain, kidney, liver, lung, and muscle tissues were fixed in 4% paraformaldehyde (PFA) in phosphate-buffered saline (PBS) before processing and embedding in paraffin. Serial 5 μm sections were mounted on glass slides. Cellular morphology was assessed using hematoxylin and eosin (H&E) staining. Cell shape, cytoplasmic integrity, and cytoskeletal organization were evaluated by immunohistochemistry using a Vimentin antibody (DAKO, Cat #M0725, 1:2,000).^25^ Apoptosis was assessed using the TUNEL assay (Millipore, Cat #S7101) with methyl green as a counterstain. Cell proliferation was evaluated using a Ki67 antibody (DAKO, Cat #M7240, 1:200).

To establish fibroblast cultures, fetal skin was digested with Collagenase Type 2 (Worthington, Cat #LS004174) in Minimum Essential Medium α (Gibco, Cat #12571-063) for approximately 2 hours at 37°C before plating in Solid Tissue Media (Minimum Essential Medium α, 10% fetal bovine serum, 1% penicillin/streptomycin, 0.2% Normocin). Astrocytes were cultured according to previously established protocols.^26^ Cell viability was estimated based on the exclusion of the vital stain, 0.4% trypan blue, prior to the initial plating of the primary cell culture.

### Nucleic Acid Extraction and Quality Analysis

Tissue samples (brain, kidney, liver, lung, muscle) designated for DNA analysis were stored at -80°C before genomic DNA extraction using the Maxwell 16 system (Promega Corporation). Agarose gel electrophoresis confirmed the presence of high molecular weight DNA. DNA integrity was assessed using an Agilent 2200 Tape Station and expressed as the DNA Integrity Number (DIN). To evaluate the suitability of genomic DNA for standard molecular diagnostic applications, polymerase chain reaction (PCR) was performed using primers for the SRY and AMELX genes: AMXY-F: CTGATGGTTGGCCTCAAGCCTGTG, AMXY-R: TAAAGAGATTCATTAACTTGACTG, SRY-F: AGTGTGAAACGGGAGAAAAC, SRY-R: TACAACCTGTTGTCCAGTTG.^27^

For RNA isolation, tissue samples (brain, kidney, liver, lung, muscle) were stored in RNAlater (Qiagen, Cat #76104) according to the manufacturer’s instructions before total RNA extraction using the Maxwell 16 system (Promega Corporation). RNA quality was assessed using an Agilent 2100 Bioanalyzer to determine the RNA Integrity Number (RIN) and the DV200 (percentage of RNA fragments >200 nucleotides).

## RESULTS

We conducted a comparative analysis of tissue samples from 18 specimens subjected to delayed evacuation (DE) after either digoxin or potassium chloride (KCl) injections, 19 to 25 hours prior, against samples from 26 specimens undergoing immediate evacuation (IE) without prior injections (Table 1 and Table S1). Among these, four specimens exhibited congenital anomalies, specifically two with hypoplastic left heart syndrome and two with various other anomalies. The average post-conception age, as indicated by fetal foot length, was significantly higher in the DE group (145 days) compared to the IE group (121 days) (Table S1). Digoxin doses ranged from 0.5 mg to 1.5 mg, with two cases receiving both digoxin and KCl. Most DE specimens displayed noticeable degradation (softening and discoloration), complicating the blinding process for analyses.

**Table 1.**
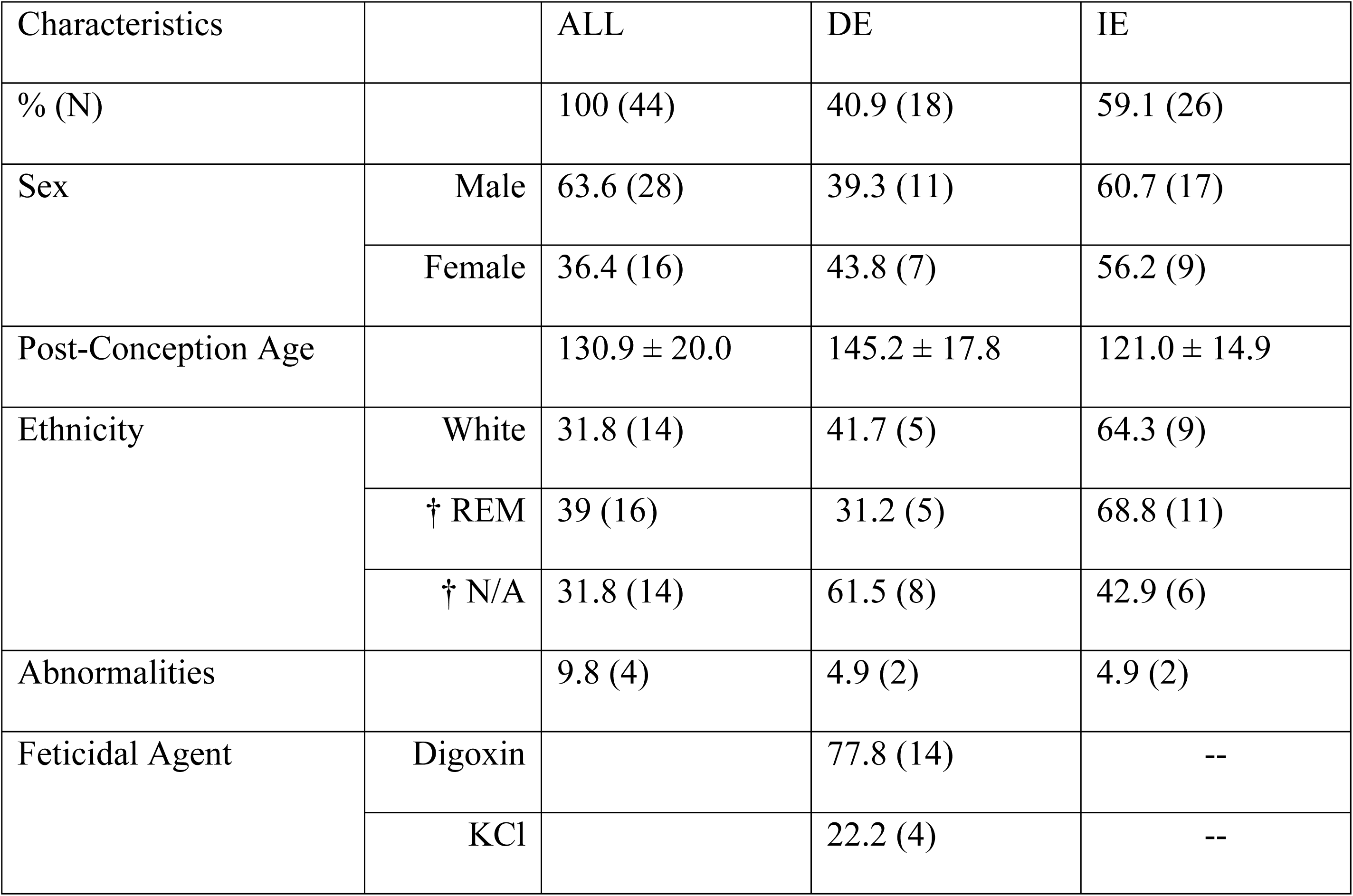
Specimen Characteristics. Characteristic summary information for the overall study population (ALL) and its stratified DE and IE subgroups. Values are presented as Mean ± SD. Welch’s t-test was used to compare age between groups, while Student’s t-test was used for other numerical variables. A p-value < 0.05 was considered statistically significant. The comparison between DE and IE for age yielded a p-value of 0.00002, indicating a significant difference. † REM defined as Racial and Ethnic Minority (excluding Hispanic); N/A defined as not available.

### Histological evaluation

We performed histological evaluations of brain, kidney, and lung from three DE specimens compared to three IE specimens (Table S1). By H&E staining, key indicators of disrupted cellular morphology including cytoplasmic swelling, vacuolization, and small, condensed nuclei that were markedly more prevalent in DE tissues (Figure 1A). Similarly, TUNEL staining was more pronounced in DE tissues, indicating severe apoptosis (Figure 1B), and Ki67 staining was nearly absent in DE tissues (Figure 1C), indicating decreased cell division and/or loss of Ki67 antigen due to degradation (Figure 1C and Table S2). Comparisons of staining estimates for both markers at 0-hour time-point can be compared by viewing DE to IE (Table S2).

**Figure 1:**
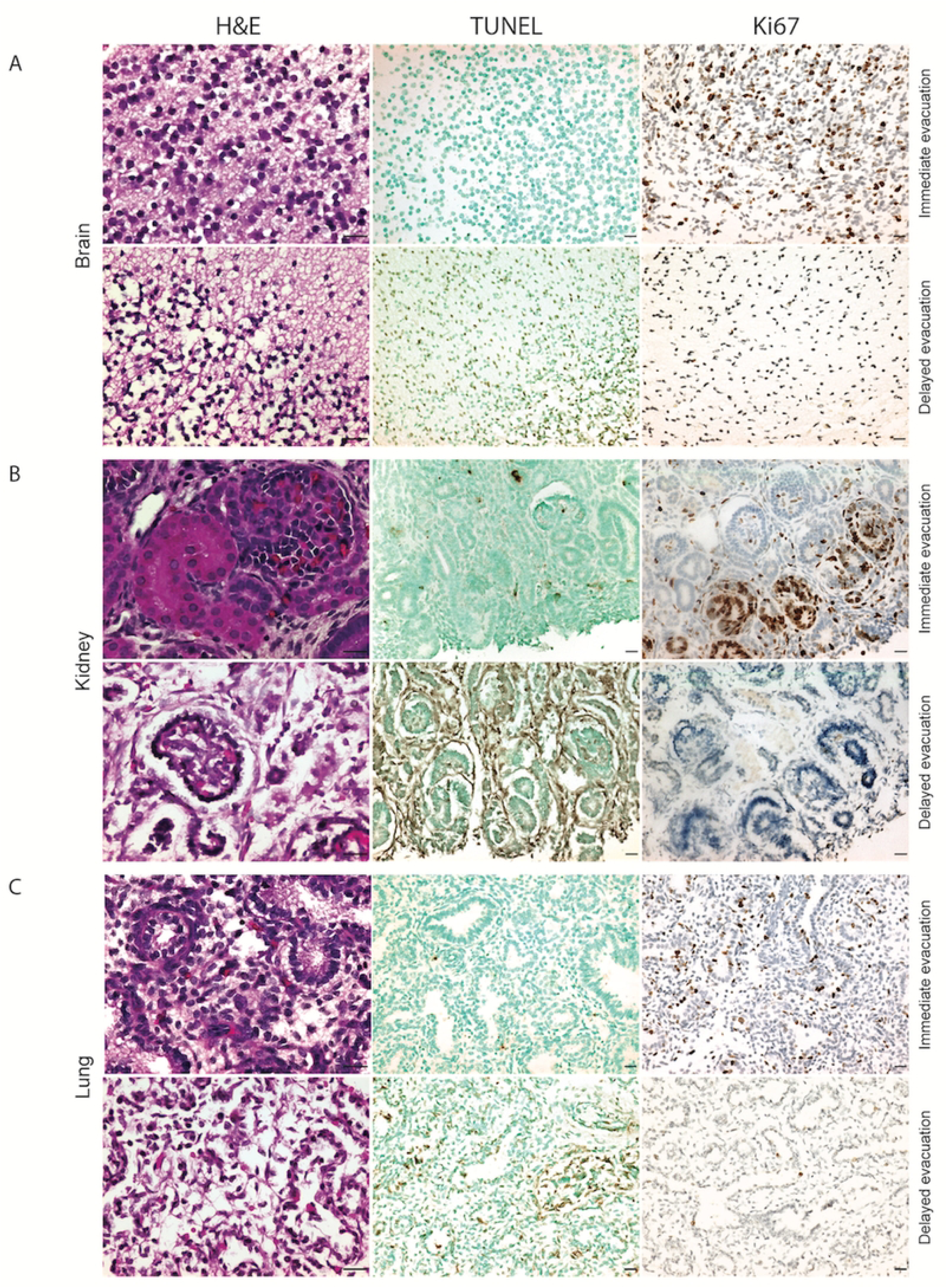

To simulate the effect of intrauterine degradation by exposure to prolonged body temperature incubation on tissue quality, we incubated IE tissues at 37°C for 6 and 18 hours (Table S2). At 6 hours, TUNEL and Ki67 staining were mildly altered compared to the 0-hour time point. At 18 hours, the incubated tissues appeared markedly degraded compared to the 0 and 6-hour time points, showed substantial TUNEL staining and lacked Ki67 staining indicating high levels of apoptosis and a complete absence of cell proliferation. The 18-hour tissues also appeared visibly more degraded than typical delayed evacuation tissues.

### Cell culture

To evaluate the ability of tissues to continue growing in culture, we sampled skin from available sites on the integument of specimens and attempted to establish fibroblast cultures. Fibroblast cultures were successfully established in 3/11 DE specimens compared to 9/9 from IE specimens. The time interval between injection of the agent and the D&E did not clearly differ between the specimens with successful and unsuccessful fibroblast cultures. In addition, the successful cultures came from specimens with and without anomalies, injected with both digoxin and KCl, and injected at different sites (intra-amniotic and intra-fetal).

We compared cell viability in three DE specimens to seven IE specimens and assayed cell viability using trypan blue staining (Figure 2). The mean percent cell viability by trypan blue staining for the DE specimens was 22% (n = 3, SD = 13%) compared to 38% (n = 7, SD = 6.7%) for IE specimens. Incubating IE brain tissue at body temperature for 6 hours lowered the viability to 16% (n = 7, SD = 5.5). Incubating for 12, 18, and 24 hours further reduced viability to 8.8% (n = 6, SD = 5.7), 5.5% (n = 6, SD = 4.5), and 1.8% (n=6, SD = 1.9) respectively. A two-sample t-test was performed to compare percent cell viability in DE tissue and IE tissue with no incubation. There was a significant difference in cell viability between DE tissue and IE tissue (p=0.033). Simple linear regression was used to test if hours incubated significantly predicted cell viability. The fitted regression model was Percent cell viability = -1.39(hours incubated) + 30.6. The overall regression was statistically significant (R² = 0.7261, F (1,30) =79.5, p<0.001).

**Figure 2:**
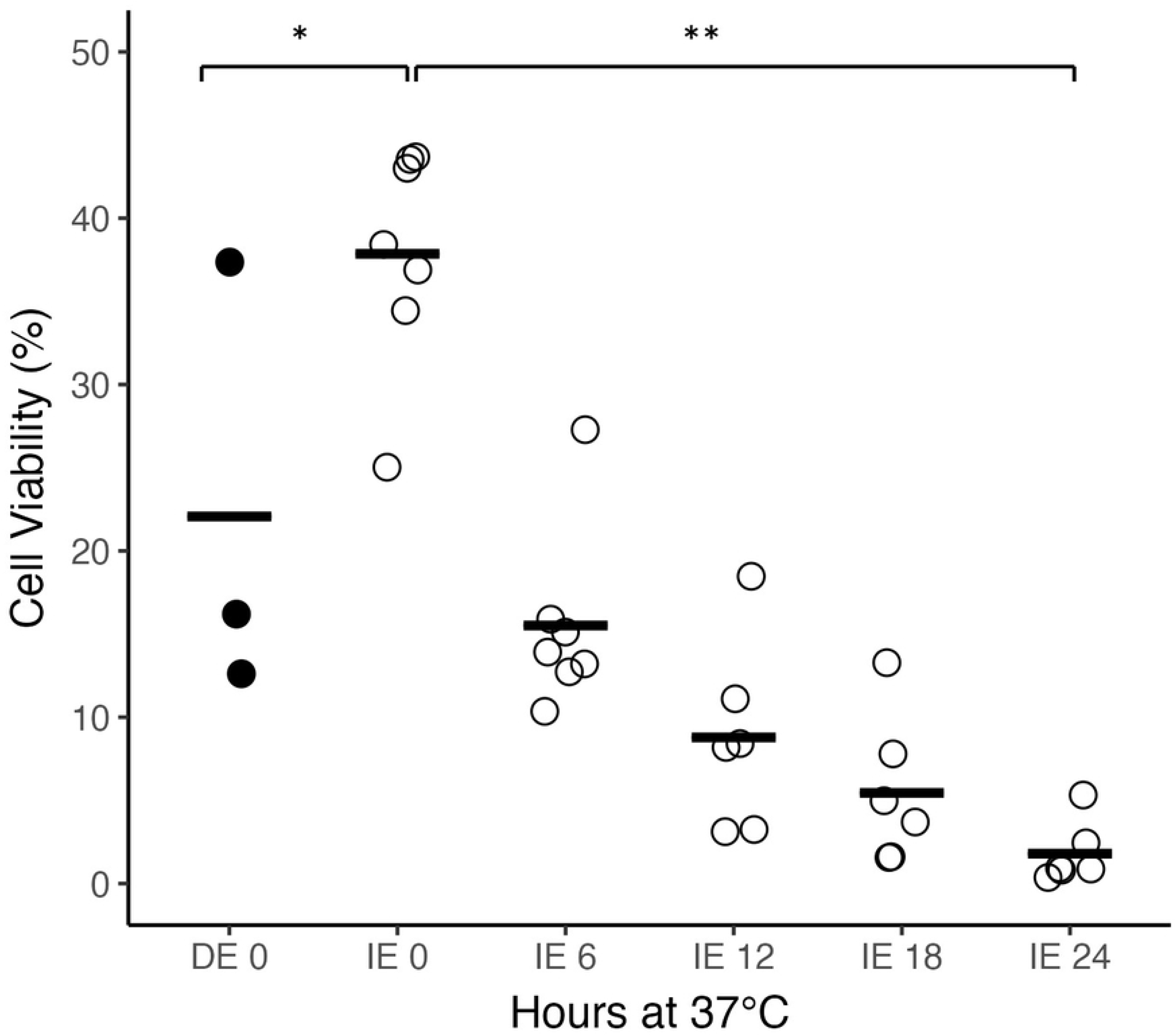

It was found that hours incubated significantly predicted cell viability (β=-1.39, p<0.001).

Despite the presence of viable cells based on trypan blue staining, no cells adhered to plates or grew from DE specimens using our standard astrocyte cell culture protocol.^26^ In contrast, astrocyte cells adhered and grew from all 11 IE brain samples. In 6/7 IE brain samples incubated for 6 hours adhered and generated cell lines. By 12 hours incubation, 2 out of the remaining 6 grew, by 18 hours 1/6 grew, and by 24 hours, no cells adhered or grew.

### Nucleic acid quality: DNA & RNA

To determine DNA quality from specimens, we isolated high molecular weight DNA from all tissues, calculated the DNA Integrity Number (DIN), and performed PCR. Based on agarose gel electrophoresis, we were able to isolate high molecular weight DNA from all tissues. However, the quality differed by organ and between DE and IE tissues. DNA extracted from brain and muscle demonstrated high molecular weight DNA with minimal smearing, whereas DNA extracted from lung, kidney, and liver showed increased smearing. While smearing may suggest DNA degradation, it can also result from factors such as RNA, protein, or salt contamination, gel preparation, well loading, or sample handling. We cannot rule out these factors entirely, though steps were taken to minimize contamination during processing.

Similarly, while DNA Integrity Numbers (DIN) were acceptable across all tissues, DE tissues generally had lower DIN scores than IE tissues (Figure S1 and Table S1). These findings indicate that while overall DNA quality remained sufficient for downstream applications such as PCR, delayed evacuation tissues may exhibit subtle compromises in DNA integrity compared to immediate evacuation tissues.

We performed PCR for sex determination using genomic DNA extracted from kidney in 8 delayed evacuation specimens and 2 immediate evacuation specimens. All 10 samples amplified appropriately, and the product sizes (977 base pairs for the X chromosome; 790 and 358 base pairs for the Y chromosome) were consistent with the external morphological sex assignment (Figure S1).

To determine the impact of DE on RNA quality, we isolated RNA from each specimen and calculated RIN scores. RIN scores for delayed evacuation tissues were all less than 6 (range 3.5-5.6) whereas Immediate evacuation tissues consistently yielded high quality RNA, with all RIN scores >8 (Figure 3A and Table S1). Using a two-sample t-test, we compared RIN scores in DE and IE tissues that were not incubated. We found significantly higher RIN scores in DE tissues for brain (p=0.0016), kidney (p<0.001), liver (p<0.001), lung (p<0.001), and muscle (p=0.0020). Next, a one-way ANOVA evaluated the effect of different incubation times (0h, 6h, 18h) on RIN scores for each organ. We observed significant changes in brain (p=0.012), kidney (p=0.0027), liver (p<0.001), lung (p=0.0011), and muscle (p<0.001). Tukey’s HSD Test showed that RIN scores generally decreased between 0h and 18h, and in many tissues between 6h and 18h as well. Brain tissue began to degrade by 6 hours at 37°C, whereas muscle was the most resistant.

**Figure 3:**
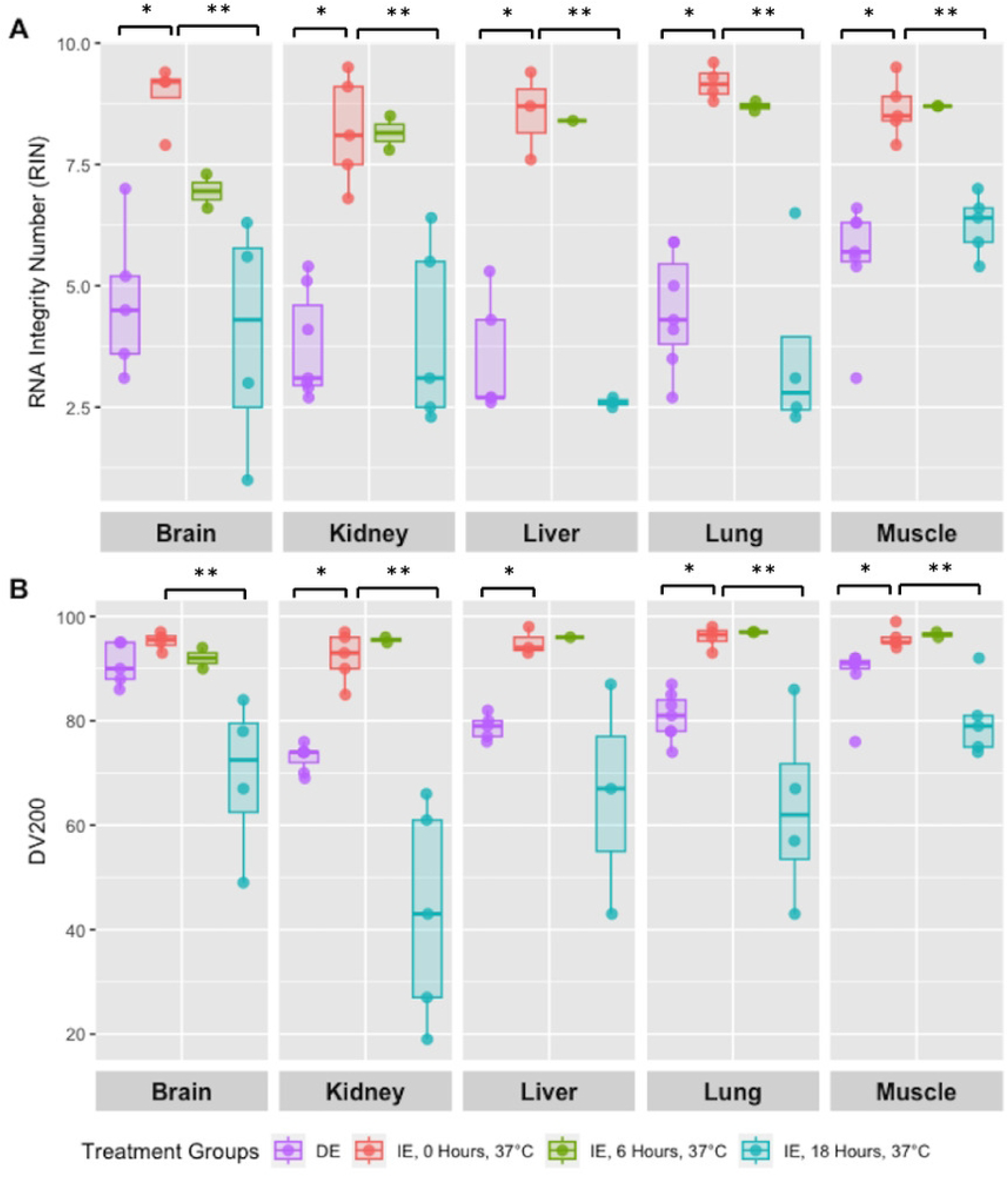

DV200 measures the percentage of RNA fragments longer than 200 nucleotides. From the recorded specimens, DV200 ranged from 73–91%, indicating moderate or better RNA quality (Figure 3B). A two-sample t-test comparing DE and IE tissues without incubation revealed significantly higher DV200 values in DE for kidney (p<0.001), liver (p=0.0010), lung (p<0.001), and muscle (p=0.019), but not for brain (p=0.073). A one-way ANOVA was then used to determine how incubation time (0h, 6h, 18h) affected DV200. Significant differences were found for brain (p=0.021), kidney (p<0.001), lung (p=0.011), and muscle (p=0.0014), but not for liver (p=0.16). Tukey’s HSD showed that DV200 values dropped between 0h and 18h in brain, kidney, lung, and muscle, and between 6h and 18h in kidney, lung, and muscle. For tissues that were not incubated, a two-sample t-test found no significant difference in RIN scores between samples with and without digoxin (brain: p=0.96, kidney: p=0.50, liver: p=0.16, lung: p=0.78, muscle: p=0.50). However, when digoxin-treated tissues were incubated for 18 hours at either 4°C or 37°C, RIN scores were significantly lower at 37°C for brain (p=0.015), kidney (p=0.0012), liver (p=0.0092), lung (p=0.025), and muscle (p=0.018).

Similarly, in non-incubated tissues, there was no significant difference in DV200 values between digoxin-treated and untreated samples (brain: p=0.76, kidney: p=0.53, liver: p=0.22, lung: p=0.76, muscle: p=0.94). But after 18 hours of incubation at 4°C or 37°C, DV200 was significantly lower at 37°C for brain (p=0.021), kidney (p=0.037), and liver (p=0.044). No significant difference was found for lung (p=0.23) or muscle (p=0.077) between these temperatures.

## DISCUSSION

### Principal Findings

While fetal pathologists commonly observe that tissues from fetuses treated with digoxin, KCl, or similar agents are often markedly autolyzed, this phenomenon has not been systematically studied. This study does not assess clinical diagnostic outcomes but instead focuses on evaluating tissue integrity under different conditions. Here, we demonstrate that induced fetal demise followed by delayed evacuation is associated with greater cellular damage and apoptosis, reduced cell proliferation, and poorer nucleic acid quality. Incubating fetal tissues at 37°C (a proxy for delayed evacuation) causes rapid degradation, while digoxin treatment alone has minimal effects regarding RNA integrity. Comparison of tissues with and without *ex vivo* digoxin exposure for 18 hours at 4°C and 37°C showed no significant effect of digoxin on RNA quality, as measured by RIN and DV200 scores (Figure 4A-4B). These findings suggest that prolonged incubation at body temperature without fetal circulation (“warm ischemia”) is the primary driver of the poorer tissue quality observed in terminated fetuses. Among the metrics assessed, cell viability and RNA integrity appear to be the most sensitive to ischemic conditions, whereas DNA integrity is relatively robust, with tissue histology and immunohistochemistry/staining falling between these extremes.

**Figure 4:**
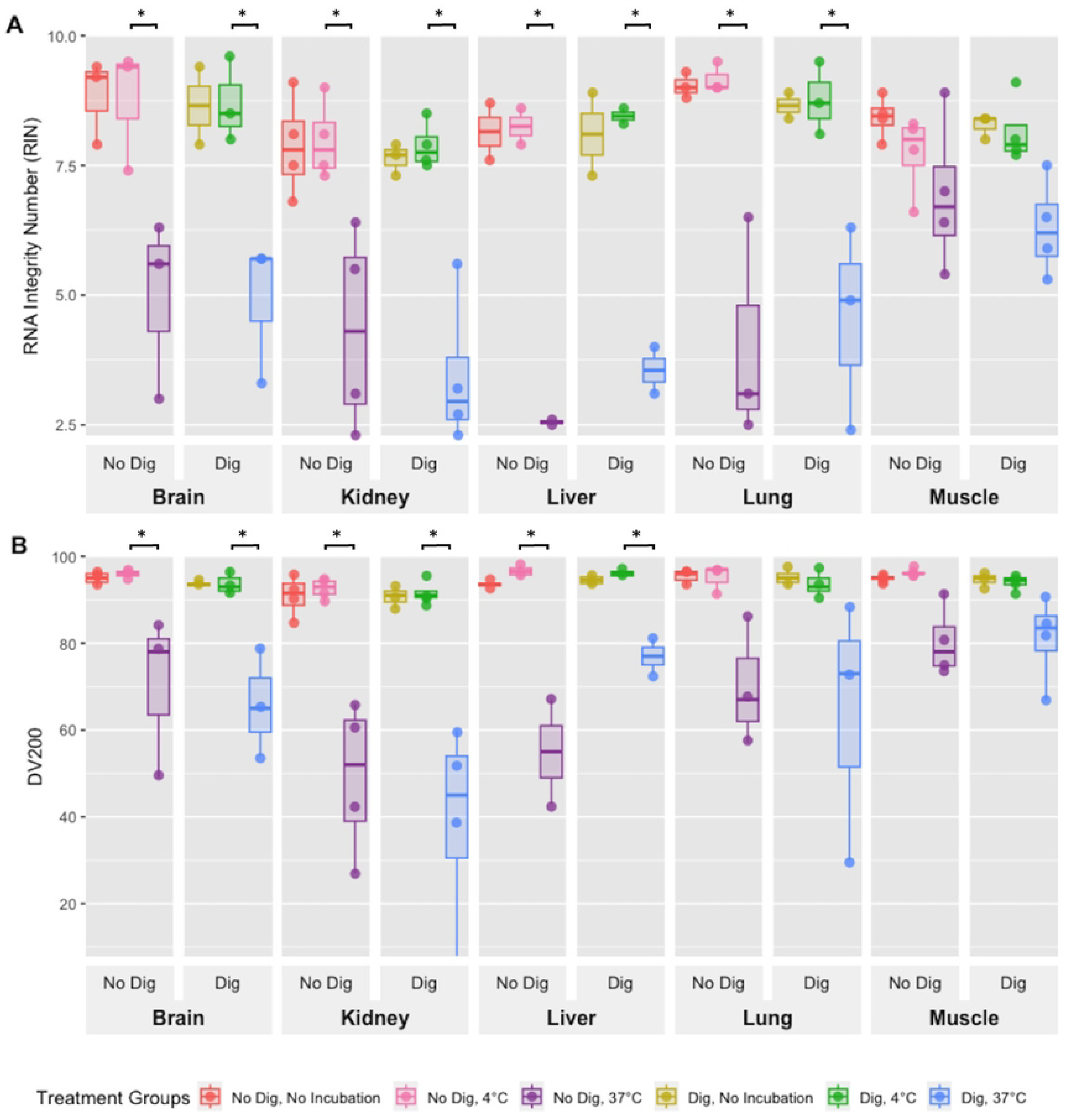

As with any medical intervention, healthcare providers and patients must weigh the risks and benefits of inducing fetal demise prior to D&E. Studies addressing the safety of digoxin for induction of fetal demise prior to termination are mixed. One study of 52 individuals randomized to two different doses of digoxin did not report adverse effects or an increase in nausea after digoxin administration.^26^ However, a large retrospective cohort study comparing patients with and without induced demise demonstrated a three-fold increased risk of complications (vomiting, intrauterine infection, and unplanned delivery).^29^ Rare but serious risks to maternal health cannot be discounted and include hyperkalemic paralysis from digoxin^30^ and cardiac arrest from KCl.^31^ No clear evidence-based medical benefits of induced fetal demise prior to D&E or induction have been reported.^21^ Patient experiences with induced fetal demise prior to D&E are complex; in one study, women reported both emotional discomfort and reassurance related to digoxin use.^32^

### Clinical and Research Implications

Different termination methods affect tissue quality in different ways. For instance, D&E causes physical disruption of the fetus, obscuring many gross anatomical features, but leaving histological features, cell viability, and nucleic acids largely intact. In contrast, induction termination results in largely intact gross anatomy. Induced fetal demise, which can occur prior to D&E or induction termination, variably impacts histology but severely impacts cell viability and RNA quality. HFT degradation affects a variety of traditional pathological evaluations, notably autopsy and its associated procedures for accurately diagnosing fetuses with abnormalities and impacting the viability of skin fibroblasts for any ancillary testing.^2^ As a result, delayed evacuation after induced demise has the potential to impair the ability of clinicians to accurately diagnose fetuses with anomalies, thereby limiting prognostic and recurrence risk counseling, and determining future reproductive options for the patient. Table S1 summarizes the outcome of the choice of termination procedure and prior induction of demise on tissue utility for both clinical and research purposes.

Although fetal DNA is relatively resistant to degradation, the increasing identification of DNA variants of uncertain significance (VUS) through clinical genetic testing often requires additional laboratory evaluations to resolve whether the variants are causal or not. For instance, DNA variants may affect RNA expression or splicing, which can be directly tested in RNA extracted from living cells, such as fibroblasts. RNA sequencing is emerging as a primary diagnostic technique, making the vulnerability of RNA especially concerning.^33,34^ Further, direct DNA sequencing of multiple affected tissues is emerging as an effective diagnostic strategy for identifying mosaic disorders, where detection of pathogenic DNA variants is restricted to a subset of tissues.^35^

Similar concerns arise for the utility of HFT in research. For many current cellular and developmental biology applications, viable cells are an essential starting material.^36^ Maintaining a mitotically active cell line, such as fibroblasts, is essential for functional work to determine how DNA variants impact RNA expression or protein translation.^37,38^ Use of HFT for research has been the focus of intense scrutiny by the general public, anti-abortion activists and the scientific community over the past few years;^37^ However, the importance of fetal tissue donation to facilitate biomedical research and health care continues to this day. In 2019, the National Institutes of Health (NIH) provided $109 million in funding for 175 studies utilizing fetal tissue for a diverse array of projects encompassing infectious disease, cancer, gene regulation, neurological disorders and degeneration, retinal disease, stem cell therapy, birth defects and the mechanisms underlying normal and abnormal fetal development.^38,39^ Two of the most notable projects, due to their scope and broad scientific impact, are BrainSpan^40^ a highly-used public resource that depicts anatomic and molecular data for the human brain across the lifespan, and ENCODE^41,42^ a large NIH-funded project that defined regulatory elements across the human genome. Ensuring that the highest quality HFT are available for research purposes also honors the intent of the tissue donation by the subject.

### Strengths and Limitations

This is the first study to systematically evaluate the impact of agents for the induction of fetal asystole on fetal tissue quality. Such agents were administered prior to D&E. We assessed quality using multiple measures relevant to the diagnosis of fetal anomalies and the use of HFT for research. This study has several important limitations, primarily related to the challenges of access to subjects and their donation. Our convenience sample made it impossible to fully match the delayed evacuation and immediate evacuation groups for fetal sex, gestational duration, and provider performing the termination. Ethically, because research participation cannot influence pregnancy decision-making or the termination method elected, we were unable to directly determine the effects of digoxin versus KCl, drug dose, injection site, or the interval between injection and D&E. Furthermore, our evaluation of cell viability and culturing capacity was restricted to fibroblasts and brain-derived cells (astrocytes). Other tissues and cell types were not tested, limiting the generalizability of our findings across all fetal tissues. Additionally, cell viability was only assessed in brain tissue, which does not capture potential variations in viability across other organ systems. These restrictions in methodology, while unavoidable due to sample availability and practical constraints, highlight the need for further research exploring tissue-specific differences in viability and culture success. Despite these limitations, the differences in tissue quality and viability observed between delayed evacuation and immediate evacuation HFT are so striking that they are unlikely to be due to confounding factors. Of note, this study does not directly address the impact of termination methods on diagnostic yield.

## CONCLUSION

Although our study did not specifically assess fetal anomalies or clinical diagnostic outcomes, it is well recognized that high-quality fetal tissue is crucial for numerous downstream analyses, including histopathology, immunohistochemistry, and molecular profiling. Our findings suggest that a shorter interval between the induction of fetal demise and evacuation is more likely to preserve tissue quality for both clinical and research applications. In settings where demise is induced prior to evacuation, providers may consider reducing this interval to ≤6 hours for patients who desire more reliable diagnostic evaluations. Intracardiac KCl or lidocaine injection, for instance, can be administered immediately before dilation and evacuation (D&E) or labor induction, facilitating expedited procedures.^19^ Additional methods to induce fetal asystole at the time of D&E include mechanical techniques, such as umbilical cord transection, which can be completed immediately prior to uterine evacuation.^21^ Ensuring that there is minimal tissue degradation can yield more promising results, enhancing the reliability of further analysis and deepening our understanding of fetal development and pathology in broader contexts.

## ACKNOWLEDGEMENTS

The authors would like to thank Raj P. Kapur, MD, PhD, from the Department of Laboratories, Seattle Children’s Hospital for his support and coordination throughout this project.

## SUPPORTING INFORMATION CAPTIONS

**Figure S1:**
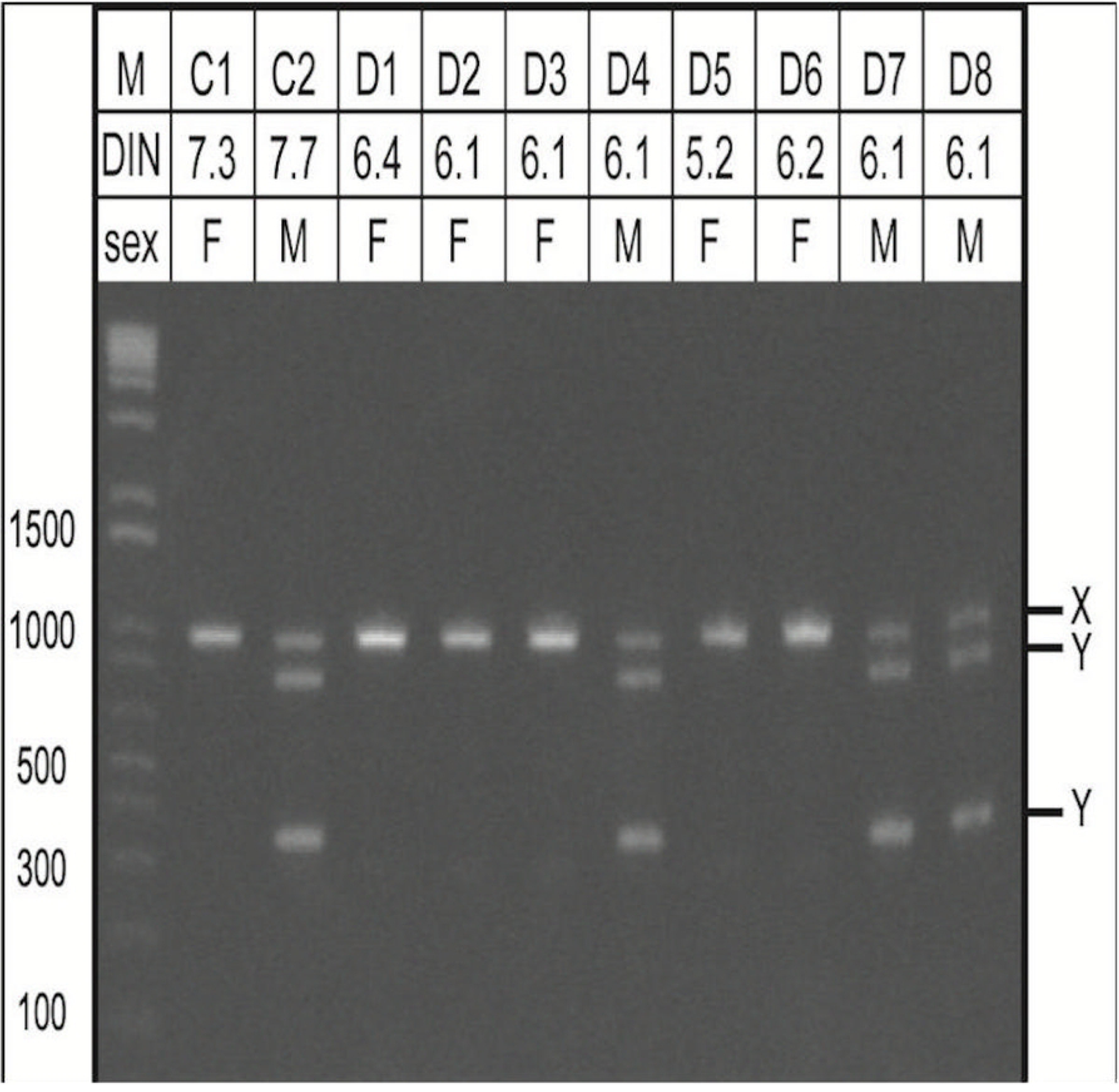
DNA Quality with Delayed Evacuation Compared to Immediate Evacuation. Sex-specific PCR products. Two IE specimens (C1-C2) and eight DE specimens (D1-D8) visualized on a 1% agarose E-Gel (Life Technologies). DNA integrity numbers (DIN) are shown above the gel. Specimens C2, D4, and D7-8 all had male external genitalia and amplified both X (977 base-pairs) and Y (790 and 358 base-pairs) chromosome-specific PCR products, while C1, D1-3, and D5-6 had female external genitalia and amplified only the X chromosome-specific PCR product. The DNA ladder (M) used is E-Gel 1Kb Plus DNA Ladder (Life Technologies #10488-090).

**Table S1: Summary of Specimen Demographics and Experimental Details.**

This table provides an overview of specimen demographics, including subject groupings (DE vs. IE), figure identifiers, DIN (DNA Integrity Number) and RIN (RNA Integrity Number) scores, DV200 scores, and the status of fibroblast culture, TUNEL, and Ki67 assays.

**Table S2: Tissue Quality Assessment by Histochemistry (TUNEL) and Immunohistochemistry (Ki67).**

Symbols indicate the proportion of cells with stain uptake compared to control tissue: (+++) 76-100%, (++) 50-75%, (+) 25-49%, (-) 0-24%. Semi-quantitative assessments of apoptosis (TUNEL) and proliferation (Ki67) in tissues from three DE specimens evacuated 19 to 25 hours after digoxin injection compared to tissues from IE specimens. Semi-quantitative assessments of apoptosis (TUNEL) and proliferation (Ki67) in tissues from two IE specimens incubated at 37°C for 0, 6, and 18 hours.

